# Comparison of the theoretical and real-world evolutionary potential of a genetic circuit

**DOI:** 10.1101/003772

**Authors:** M. Razo-Mejia, J. Q. Boedicker, D. Jones, A. DeLuna, J. B. Kinney, R. Phillips

**Affiliations:** Ingenieria Biotecnologica, Instituto Politecnico Nacional, Av. Mineral de Valenciana No. 200 Col. Fracc. Industrial Puerto Interior, Silao de la Victoria, Guanajuato, 36275, Mexico.; Department of Applied Physics, California Institute of Technology, 1200 East California Boulevard, Pasadena, California 91125, USA.; Laboratorio Nacional de Genomica para la Biodiversidad (Langebio), Centro de Investigacion y de Estudios Avanzados (Cinvestav), 36821 Irapuato, Guanajuato, Mexico.; Simons Center for Quantitative Biology, Cold Spring Harbor Laboratory, Cold Spring Harbor, NY 11724

**Keywords:** thermodynamic models, *lac* operon, evolutionary potential, transcriptional regulation, natural variability

## Abstract

With the development of next-generation sequencing technologies, many large scale experimental efforts aim to map genotypic variability among individuals. This natural variability in populations fuels many fundamental biological processes, ranging from evolutionary adaptation and speciation to the spread of genetic diseases and drug resistance. An interesting and important component of this variability is present within the regulatory regions of genes. As these regions evolve, accumulated mutations lead to modulation of gene expression, which may have consequences for the phenotype. A simple model system where the link between genetic variability, gene regulation and function can be studied in detail is missing. In this article we develop a model to explore how the sequence of the wild-type *lac* promoter dictates the fold change in gene expression. The model combines single-base pair resolution maps of transcription factor and RNA polymerase binding energies with a comprehensive thermodynamic model of gene regulation. The model was validated by predicting and then measuring the variability of *lac* operon regulation in a collection of natural isolates. We then implement the model to analyze the sensitivity of the promoter sequence to the regulatory output, and predict the potential for regulation to evolve due to point mutations in the promoter region.

## 1. Introduction

Despite efforts to understand genotypic variability within natural populations [1] and recent interest in fine-tuning genetic circuits for synthetic biology [2], it still remains unclear how, with base pair resolution, the sequence of a gene regulatory region can be translated into output levels of gene expression [3]. Generally, classical population genetics has treated regulatory architectures as changeless parameters, rather than potential evolutionary variables, focusing on changes in protein structure rather than gene regulation. However, genetic regulatory architecture can also determine the variation of traits, and thus the evolutionary potential of these genes [4]. After all, the structure of bacterial promoters dictates interactions among the transcriptional apparatus, and through the modification of this structure, regulatory circuits can be modified to potentially allow cells to occupy different niches [5, 6].

Thermodynamic models of gene regulation have been widely used as a theoretical framework to dissect and understand genetic architectures [7, 8, 9, 10, 11]. Such dissections have led to a quantitative understanding of how parameters such as binding energies, transcription factor copy numbers, and the mechanical properties of the DNA dictate expression levels. Recently the development of experimental techniques combining these types of models with cell sorting and high-throughput sequencing have made it possible to understand gene regulation at single-base pair resolution [12, 13, 14], as well as to deliberately design promoter architectures with desired input-output functions [15]. These models connect the sequence of a promoter to the output phenotype, making it possible to predict variability and evolutionary potential of gene regulatory circuits.

The *lac* operon has served as a paradigm of a genetic regulatory system for more than 60 years [16, 17]. This operon contains the molecular machinery that some bacterial species, including the model organism *E. coli*, use to import and consume lactose. Extensive quantitative characterization of the regulation of this genetic circuit [18, 19], as well as of the link between fitness and expression of the operon [20, 21, 22, 23, 24] make it an ideal system for exploring the evolutionary potential of a regulatory circuit. With previous exhaustive description and quantification of the parameters controlling the expression level of this genetic circuit [19, 25, 26, 27] we now have what we think is a nearly complete picture of the regulatory *knobs* that can modify the expression level, shown schematically in Figure 1(a). In this article we build upon this understanding by directly linking the sequence of the promoter region with these control parameters, thereby creating a map from genotype to transcriptional output.

**Figure 1.**
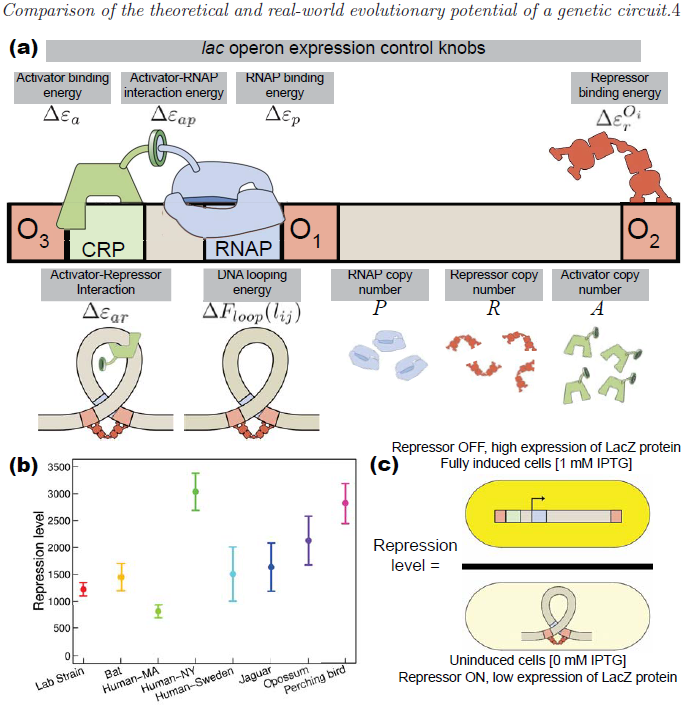
(a) Regulatory knobs that control the expression of the *lac* operon and the symbols used to characterize these knobs in the thermodynamic model. The activator CRP increases expression, the Lac repressor binds to the three operators to decreases expression, and looping can lock the repressor onto *O*_1_ leading to increased repression. The interaction energy between RNAP and CRP reflects the stabilization of the open complex formation due to the presence of the activator [28], and the interaction between the Lac repressor and CRP stabilizes the formation of the upstream loop [29]. (b) Variability in the repression level of *E. coli* natural isolates and the lab control strain MG1655. Strains are named after the host organism from which they were originally isolated [30]. Error bars represent the standard deviation from at least three independent measurements. (c) Schematic representation of the repression level, in which the role of the repressor in gene regulation is experimentally measured by comparing the ratio of LacZ proteins in cells grown in the presence of 1 mM IPTG to cells grown in the absence of IPTG. LacZ protein concentrations were measured using a colorimetric assay.

Within a collection of *E. coli* isolated from different host organisms we observe significant variability for the regulation of the *lac* operon, as shown in Figure 1(b). By characterizing the variability of the regulatory control parameters shown in Figure 1(a) within these strains, we identified evolutionary trends in which certain parameters or subsets of parameters are seen to vary more often than others within this collection of natural isolates. Using the map of promoter sequence to transcriptional output, we demonstrated that the regulatory input-output function for the *lac* promoter could account for most of the natural variability in regulation we observed. We then implement the map to explore the theoretical potential for this regulatory region to evolve. This level of analysis gives us clues as to how selection could fine tune gene expression levels according to the environmental conditions to which cells are exposed.

## 2. Results

### 2.1. Quantitative model of the natural parameters that regulate gene expression

Thermodynamic models of gene regulation have become a widely used theoretical tool to understand and dissect different regulatory architectures [3, 12, 19, 26, 27, 31]. The *lac* promoter is one such regulatory architecture that has been studied in detail [32]. Models have been constructed and experimentally validated for both the wild-type *lac* promoter and synthetic promoter regions built up from the *lac* operon’s regulatory components [12, 15, 19, 26, 27, 32, 33, 34, 35, 36, 37]

In a simple dynamical model of transcription the number of messenger RNA (mRNA) is proportional to the transcription rate and the degradation rate of the mRNA,

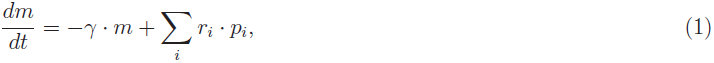

where *γ* is the mRNA degradation rate and *m* is the number of transcripts of the gene per cell; *r*_*i*_ and *p*_*i*_ are the transcription rate and the probability of state *i* respectively. We can think of *p*_*i*_ as a measure of the time spent in the different transcriptionally active states. Thermodynamic models assume that the gene expression level is dictated by the probability of finding the RNA *polymerase* (RNAP) bound to the promoter region of interest [7, 8, 9]. With a further quasi-equilibrium assumption for the relevant processes leading to transcription initiation, we derive a statistical mechanics description of how parameters such as transcription factor copy number and their relevant binding energies, encoded in the DNA binding site sequence, affect this probability [10]. Quantitative experimental tests of predictions derived from equilibrium models have suggested the reasonableness of the assumption [15, 19, 26, 27], although caution should be used as the equilibrium assumption is not necessarily valid in all cases. The validity of this equilibrium assumption relies on the different time-scales of the processes involved in the transcription of a gene. Specifically the rate of binding and unbinding of the transcription factors and the RNAP from the promoter region should be faster than the open complex formation rate; if so, the probability of finding the RNAP bound to the promoter is given by its equilibrium value [9, 38]. For the case of the Lac repressor, the rate of unbinding from the operator is 0.022 1/s [39], and the binding of an unoccupied operator with 10 repressors per cell occurs at a similar rate [40]. Open complex formation, a rate limiting step in promoter escape, has been measured at a rate of 2 *×* 10^−3^ 1/s [41]. Promoter escape is about an order to magnitude slower than the binding and unbinding of the Lac repressor, and this separation of time scales supports the equilibrium assumption for this particular case. We enumerate the possible states of the system and assign statistical weights according to the Boltzmann distribution as shown in Figure 2.

**Figure 2.**
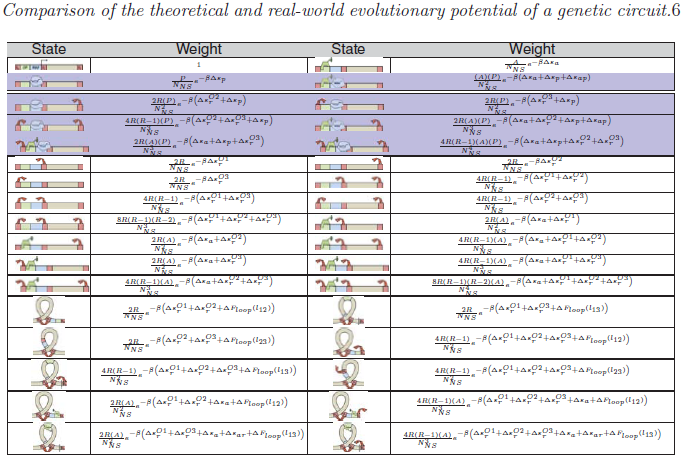
Thermodynamic model of gene regulation. The table shows all states permitted within the model and their respective statistical weights as obtained using statistical mechanics. In these weights *P* = number of RNAP per cell, *R* = number of repressor molecules per cell, *A* = number of activator molecules per cell, 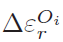 = binding energy of Lac repressor to the *i^th^* operator, Δ*ε_p_* = binding energy of RNA polymerase to the promoter, Δ*ε_a_* = activator binding energy, Δ*F*_*loop*_(*l*_*ij*_) = looping free energy between operator *O*_*i*_ and *O*_*j*_, *N*_*NS*_ = number of nonspecific binding sites on the genome, Δ*ε_ap_*= interaction energy between the activator and the RNAP, Δ*ε_ar_*= interaction energy between the activator and the repressor, and *β* = inverse of the Boltzmann constant times the temperature (see Supplemental Material for further discussion). States with blue background are assumed to lead to transcription of the operon.

From these states and weights we derive an equation describing the probability of finding the system in a transcriptionally active state, and therefore the production term from Equation 1,

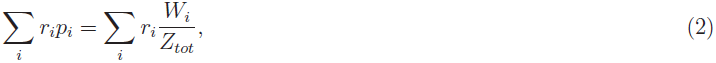

where *W*_*i*_ is the statistical weight of states in which the polymerase is bound, which are assumed to lead to the transcription of the operon (shaded blue in Figure 2), and *Z*_*tot*_ = Σ_All states_ *W*_*state*_ is the partition function, or the sum of the statistical weights of all states. We connect this model to experimental measurements of repression, that is the ratio of gene expression in the absence of the active repressor to gene expression in the presence of active repressor, using:

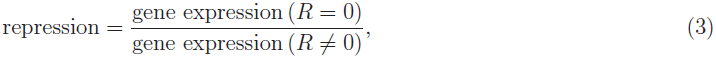

where *R* is the number of repressor molecules per cell. The experimental equivalent of repression is depicted in Figure 1(c). In experiments, isopropyl *β*-D-1-thiogalactopyranoside (IPTG) is used to inactivate the Lac repressor, preventing it from binding to the genome with high affinity [19]. Repression, as defined in Equation 3, has been a standard metric for the role of transcription factors, including the Lac repressor, on gene expression [7, 42]. By measuring the ratio of steady-state levels of a gene reporter protein, here LacZ, we are able to isolate the role of the repressor in gene regulation, as described further in section S8 of the Supplemental Material.

Various models of the wild-type *lac* promoter have been reported in the past using this simple structure. Our work builds upon the work by Kinney *et al.* [12]. Kinney and collaborators combined a thermodynamic model of regulation with high-throughput sequencing to predict gene expression from statistical sequence information of the cAMP-receptor protein (CRP) and the RNAP binding sites. To predict how the sequence of the entire regulatory region influences expression, we adapted this model to account for how the binding site sequence and copy number of the Lac repressor modulate gene expression. Our model also takes into account growth rate effects, captured in the RNAP copy number [43, 44].

Based on previous work done on the *lac* operon [19, 12], we assumed that the presence of the activator does not affect the rate of transcription (*r*_*i*_ from Equation 1), but instead influences the probability of recruiting the polymerase to the promoter (*p*_*i*_ from Equation 1). Previous experimental characterization of the repressor binding energy to the different operators [26], the looping free energy for the upstream loop between *O*_1_ – *O*_3_ [27], activator concentration and its interaction energy with RNAP [19], RNAP binding energy [15] and RNAP copy number as a function of the growth rate [44], left us only with three unknown parameters for the model. One of these missing parameters, a decrease in the looping free energy when CRP and Lac repressor are bound at the same time, is a consequence of the experimental observation that the presence of CRP stabilizes the formation of the loop between *O*_1_ – *O*_3_ [29, 45]. The remaining two parameters, the looping energies for the *O*_1_*–O*_2_ and *O*_3_*–O*_2_ loops are not well characterized. These looping energies may differ from upstream loops due to the absence of the RNAP binding site which modifies the mechanical properties of the loop [46]. We fit these parameters for our model using Oehler *et al.* repression measurements on *lac* operon constructs with partially mutagenized or swapped binding sites [42, 47] (see section S5 of the Supplemental Material for further details). Using these parameters the model is consistent with previous measurements (Figure S4). We emphasize that having the 14 parameters of the model characterized (see Table S1) provides testable predictions without free parameters that we compare with our experimental results.

### 2.2. Sensitivity of expression to model parameters

As an exploratory tool, the model can predict the change in regulation due to modifications in the promoter architecture. Figure 3 shows the fold-change in the repression level as a function of each of the parameters, using the lab strain MG1655 as a reference state (see Supplemental Material for further detail on these reference parameters). We have reported parameters using strain MG1655 as a reference strain because this strain served as the basis for which most parameter values were determined and the gene expression model was derived.

**Figure 3.**
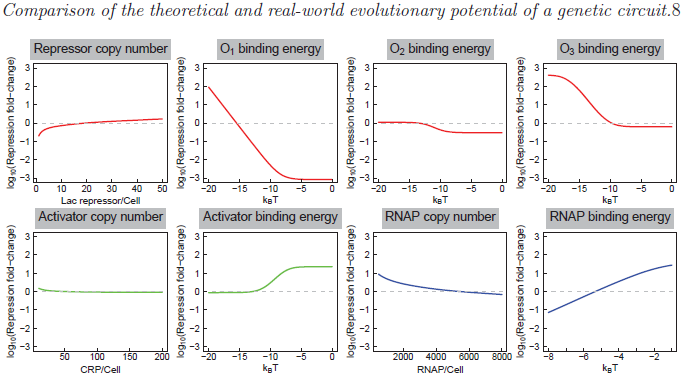
Sensitivity of phenotype to the parameters controlling the gene expression level. Each graph shows how a specific model parameter changes the level of gene expression. The log_10_ ratio of repression is calculated with respect to the predicted repression for the lab strain MG1655. The vertical axis spans between 1000 fold decrease to 1000 fold increase in repression with respect to this strain. The gray dotted line indicates the reference value for the lab strain MG1655. Values above this line indicate the operon is more tightly repressed and values below this line have a leakier expression profile (see Table S1 for further detail on the reference parameters).

From this figure we see that within the confines of this model, modifications in the *O*_1_ binding energy have the most drastic effect on the repression of the operon. For the case of *O*_2_ we see that increasing its affinity for the repressor does not translate into an increased ability to turn off the operon; but by decreasing this operator affinity the model predicts a reduction in the repression with respect to the reference strain.

Surprisingly the repression level is predicted to be insensitive to activator copy number. The same cannot be said about the affinity of the activator, since decreasing the activator binding energy greatly influences the repression level.

### 2.3. Mapping from sequence space to level of regulation

Recent developments of an experimental technique called *sort-seq*, involving cell sorting and high-throughput sequencing, have proved to be very successful in revealing how regulatory information is encoded in the genome with base pair resolution [12]. This technique generates energy matrices that make it possible to map from a given binding site sequence to its corresponding binding energy for a collection of different proteins and binding sites. Combining these energy matrices with thermodynamic models enables us to convert promoter sequence to the output level of gene expression. Recently these energy matrices have been used to deliberately design promoters with a desired expression level, demonstrating the validity of these matrices as a design tool for synthetic constructs [15]. We use the matrices for CRP and RNAP published previously [12]. We experimentally determined the matrix for the LacI operator using previously published methods [12], as discussed in Materials and Methods. Figure 4(a) shows a schematic representation of the relevant protein binding sites involved in the regulation of the *lac* operon and their respective energy matrices. Implementing these matrices into the thermodynamic model gives us a map from genotype to phenotype. We use this map to calculate the fold-change in repression relative to MG1655 for all possible point mutations in this region. Figure 4(b) shows the fold-changes in repression levels for the two base pair substitutions at each position that result in the largest predicted increase or decrease in repression.

**Figure 4.**
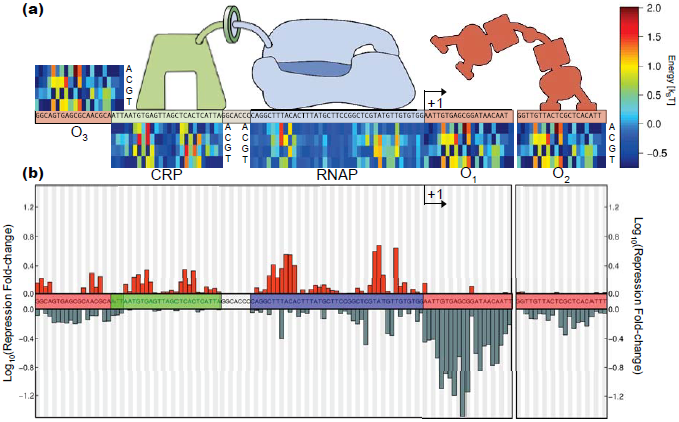
Mapping from promoter sequence to regulatory level. (a) Energy matrices for the relevant transcription factors (Blue - RNAP, green - CRP, red - Lac repressor). These matrices allow us to map from sequence space to the corresponding binding energy. The contribution of each base pair to the total binding energy is color coded. The total binding energy for a given sequence is obtained by adding together the contribution of each individual base pair. (b) Using the energy matrices from (a) and the model whose states are depicted in Figure 2, the log_10_ repression change was calculated for all possible single point mutations of the promoter region. The height of the bars represents the biggest possible changes in the repression level (gray bars for biggest predicted decrease in repression, orange bar for biggest predicted increase in repression) given that the corresponding base pair is mutated with respect to the reference sequence (*lac* promoter region of the lab strain MG1655). The black arrows indicate the transcription start site.

Again we see that mutations in the *O*_1_ binding site have the largest effect on regulation since a single base pair change can lower the ability of the cell to repress the operon by a factor of *≈* 20. With only two relevant mutation that could significantly increase the repression level, this map reveals how this operator and its corresponding transcription factor diverged in a coordinated fashion; the wild-type sequence has nearly maximum affinity for the repressor [48]. It is known that the non-natural operator *O*_*id*_ binds more strongly than *O*_1_ [42]. *O*_*id*_ is one base pair shorter than *O*_1_ and current maps made with *sort-seq* cannot predict changes in binding affinity for binding sites of differing length, although accounting for length differences in binding sites is not a fundamental limitation of this method.

For the auxiliary binding sites, the effect discussed in section 2.2 is reflected in this map: increasing the Lac repressor affinity for the *O*_2_ binding site does not increase repression. Mutations in almost all positions can decrease repression, and no base pair substitutions significantly increase the repression level. Mutations in the *O*_3_ binding site have the potential to either increase or decrease the repression level. With respect to the RNAP binding site, we can see that, as expected, the most influential base pairs surround the well characterized -35 and -10 boxes. The CRP binding site overlaps three base pairs with the upstream Lac repressor auxiliary operator. As the heat-map reveals, the binding energy is relatively insensitive to changes in those base pairs, so we assume independence when calculating the binding energy and capture the synergy between the Lac repressor bound to *O*_3_ and CRP with an interaction energy term.

The construction of the sequence to phenotype map enables us to predict the evolvability of the *lac* promoter region. We calculated the effect that all possible double mutations would have in the regulation of the operon, again with respect to the predicted repression level of the reference strain MG1655. Figure 5 shows what we call the “phenotype change distribution” obtained by mutating one or two base pairs from the reference sequence, under the assumption of same growth rate and transcription factor copy numbers as the reference strain. The distribution peaks at zero for both cases, meaning that the majority of mutations are predicted not to change the repression level with respect to the reference strain, and would result in genetic drift. However it is interesting to note that the range of repression values predicted by the model with only one mutation varied between 30 times lower and 4.6 times higher than the reference value, and with two mutations the repression varied between 345 times lower and 15 times higher than the reference value. This suggests that regulation of this operon could rapidly adapt and fine tune regulation given appropriate selection.

**Figure 5.**
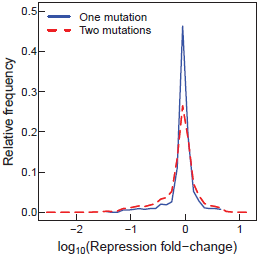
Phenotype change distribution. Relative frequency of the predicted changes in repression level by mutating one (solid blue line) or two (dashed red line) base pairs from the reference sequence (MG1655 promoter region).

### 2.4. Promoter sequence variability of natural isolates and available sequenced genomes

In order to explore the natural variability of this regulatory circuit, we analyzed the *lac* promoter region of 22 wild-type *E. coli* strains which were isolated from different organisms [30], along with 69 fully sequenced *E. coli* strains (including MG1655) available online (http://www.ncbi.nlm.nih.gov/genomes/MICROBES/microbial_taxtree.html). Figure 6 summarizes the sequencing results; for comparison, we plot the “genotype to phenotype map” from Figure 4(b) to gain insight into how the sequence variability influences regulation in these strains. Figure 6(b) shows the relative frequency of single nucleotide polymorphisms (SNP’s) with respect to the consensus sequence. Qualitatively we can appreciate that the mutations found in these strains fell mostly within base pairs which according to the model weakly regulated expression. To quantify this observation we mapped the sequences to their corresponding binding energies. As shown in Figure 6(c) the distribution of parameters is such that the observed mutations result in relatively small changes to the binding energies, less than 1 *k_B_T* relative to the reference sequence, except for the *O*_3_ binding energy that is predicted to increase *>*1 *k_B_T* in 16 strains.

**Figure 6.**
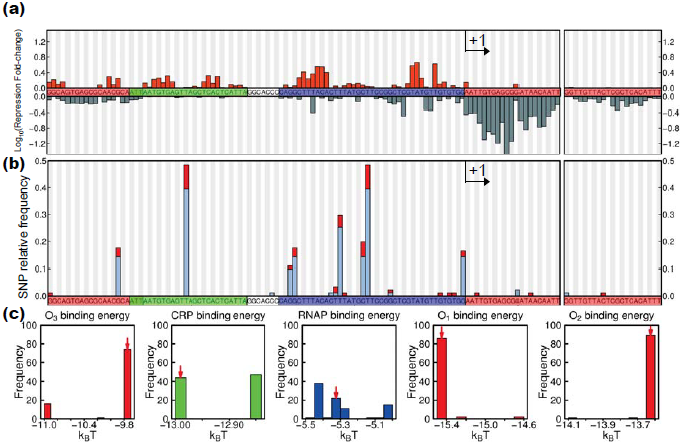
Mutational landscape of the regulatory region of the *lac* operon. (a) The genotype to phenotype map is reproduced from Figure 4(b) in order to show how each base pair in the region influences gene regulation. (b) Comparing the sequence of the *lac* promoter from 91 *E. coli* strains identifies which base pairs were mutated in this region. The height of the bars represent the relative frequency of a mutation with respect to the consensus sequence. The red part of each bar represents the 22 natural isolates from different hosts [30] and the light blue part of these bars represents the 69 fully sequenced genomes (http://www.ncbi.nlm.nih.gov/genomes/MICROBES/microbial_taxtree.html). Color coding of the binding sites and the transcription start site is as in Figure 4. (c) Using the energy matrices of Figure 4(a), we calculate the variability of protein binding energies for all sequences. The red arrow indicates reference binding energies for control strain MG1655.

### 2.5. Does the model account for variability in the natural isolates?

Next we further characterized the eight strains from Figure 1(b) in order to determine if the observed variability in regulation could be accounted for in the model (see section S2 for details on the 16S rRNA of this subset of strains). In particular, we measured the *in vivo* repressor copy number with quantitative immunoblots (see Material and Methods) and the growth rate. Table 1 shows the measured repressor copy number and the doubling time for these strains.

**Table 1.**
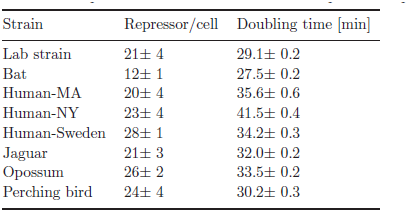
Lac repressor copy number as measured with the immunodot blots and doubling time of the eight strains with measured repression level shown in Figure 1(b). The errors represent the standard error of 3 independent experiments.

| Strain | Repressor/cell | Doubling time [min] |
| --- | --- | --- |
| Lab strain | $21 \pm 4$ | $29.1 \pm 0.2$ |
| Bat | $12 \pm 1$ | $27.5 \pm 0.2$ |
| Human-MA | $20 \pm 4$ | $35.6 \pm 0.6$ |
| Human-NY | $23 \pm 4$ | $41.5 \pm 0.4$ |
| Human-Sweden | $28 \pm 1$ | $34.2 \pm 0.3$ |
| Jaguar | $21 \pm 3$ | $32.0 \pm 0.2$ |
| Opossum | $26 \pm 2$ | $33.5 \pm 0.2$ |
| Perching bird | $24 \pm 4$ | $30.2 \pm 0.3$ |

Using the thermodynamic model by taking into account the repressor copy number, the promoter sequence and the growth rate, we predict the repression level for each of the isolates measured in Figure 1(b). In Figure 7 we plot these predicted values vs. the experimental measurements. We find that the model accounts for the overall trends observed in the isolates, with the predictions for 6 of 8 strains falling within two standard deviations of the measurements. A few of the measured repression values fall outside of the prediction, suggesting that the model may not capture the full set of control parameters operating in all of the strains.

**Figure 7.**
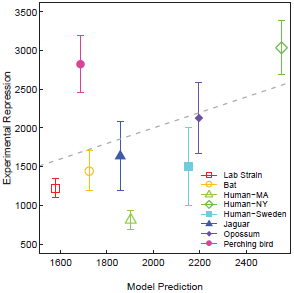
Comparison of model predictions with experimental measurements. Error bars represent the standard deviation of at least 3 independent measurements each with three replicates. The dotted line plots *x* = *y*.

### 2.6. Exploring the variability among different species

We extended our analysis to different microbial species with similar *lac* promoter architectures. After identifying bacterial species containing the *lac* repressor, we used the *sort-seq* derived energy matrices shown in Figure 4(a) to identify the positions of the transcription factor binding sites in each of these candidate strains. We identified a set of eight species whose *lac* promoter architecture was similar to *E. coli*. Figure 8 shows the 16S rRNA phylogenetic tree for these strains. The predicted change in regulation was calculated for these strains using the model whose states are shown in Figure 2, the energy matrices in Figure 4(a), and assuming all strains have the same growth rate and transcription factor copy numbers as the lab strain MG1655. The repression level relative to *E. coli* among these species is predicted to increase as much as a factor of *≈* 20 and decrease as much as a factor of *≈* 4. Regulation of the operon seems to follow phylogenetic patterns in the 16S rRNA tree, with *E. coli* relatives having a similar predicted repression level, *Citrobacter* evolved to increase repression, and *Salmonella* evolved to decrease repression.

**Figure 8.**
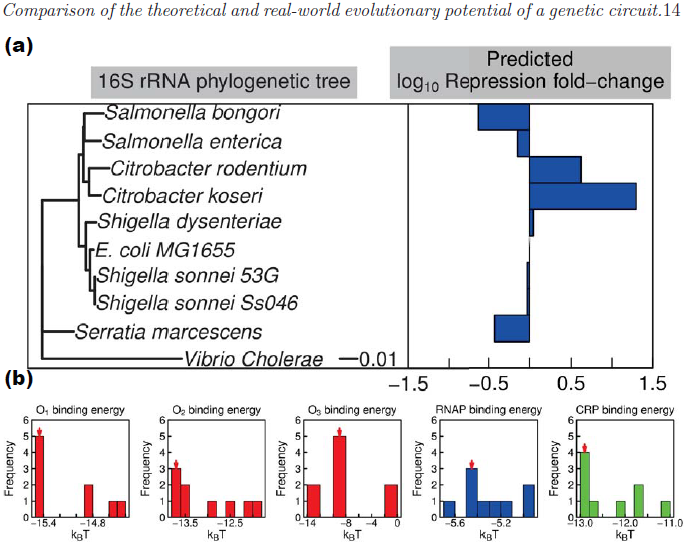
Predicted variability among different microbial species based on genome sequences and our model for regulation derived for *E. coli*. (a) On the left a 16S rRNA phylogenetic tree of diverse species with a similar *lac* promoter architecture done with the Neighbor-Joining algorithm. *Vibrio cholerae* was used as an outgroup species. The scale bar represents the relative number of substitutions per sequence. On the right the predicted log_10_ fold-change in repression with respect to *E. coli* MG1655 assuming the same growth rate and transcription factor copy numbers. The outgroup species fold-change was not calculated. (b) Parameter distribution calculated using the promoter region sequence and the energy matrices. The red arrow indicates the MG1655 reference value. Strains lacking a binding site were binned as zero.

## 3. Discussion

The approach presented here combines thermodynamic models of gene regulation with energy matrices generated with *sort-seq* to produce a single-base pair resolution picture of the role that each position of the promoter region has in regulation. These types of models based on equilibrium statistical mechanics have been used previously for the *lac* operon [19, 25], here we expanded the model to account for important cellular parameters such as growth rate, the binding site strengths of all transcription factors, and the binding site strength of RNAP. Thermodynamic models are functions of the natural variables of the system as opposed to the widely used phenomenological Hill functions [49], where it is less straightforward how changes to a promoter region translate to changes in regulatory parameters such as *K*_*M*_, the half saturation constant, and *n*, the Hill coefficient. Currently our model assumes that protein-protein interactions and DNA looping energies are kept constant, but these variables could also be a function of the promoter sequence, affecting the positioning of the transcription factors and therefore their interactions with the other molecules involved.

The underlying framework developed here can be applied to any type of architecture. Here we use the *lac* operon because it is well characterized. There is no reason to believe that this approach could not be extended to other regulatory regions, however such an effort would require extensive quantitative characterization of the control parameters of each genetic circuit, such as protein copy numbers, interaction energies, and binding affinities. Although this level of characterization requires additional experimental effort, we believe that developing such predictive, single-base pair models of gene regulation can lead to significant insights into how genetic circuits function, interact with each other, and evolve.

The majority of the natural variability found among the sequenced promoters tended to fall in bases predicted to have low impact on overall regulation, as shown in Figure 6. As an example the highly conserved mutation in the CRP binding energy or the mutations along the RNAP binding site are predicted to change the binding energy by less than 1 *k_B_T*, having a very low impact on the repression level. With respect to the repressor binding sites, among the sequenced natural isolates only one mutation was found in the *O*_2_ binding site. Unlike the *O*_1_ and *O*_3_ operators, the evolution of *O*_2_ may be constrained given that its sequence encodes both gene regulatory information and is part of the coding region of the *β*-galactosidase gene.

As shown in Figure 7, after taking into account the variability in the promoter sequence, changes in the repressor copy number, and changes in the growth rate the model accounts for most of the variability in regulation for the majority of the isolates. Linear regression of the entire experimental dataset weighted by the inverse of their standard deviation gives a slope of 1.26 with an *R*^2^ of 0.24. It can be seen that many of the points fall close to or on the x=y line, indicating that the poor fit is a result of a few outliers within the dataset. Removing the outliers (Perching bird, Human-MA, and Human-NY) results in a best fit line of slope 1.05 with *R*^2^ 0.74, reiterating that the model is consistent with the phenotype of 5 of 8 isolates. It is interesting that the three isolates whose regulatory outputs were predicted poorly by the model (Perching bird, Human-MA, and Human-NY in Figure 7) all have identical promoter sequences, which is the consensus promoter sequence as shown in Figure S1. Although these three strains have identical sequences, two strains repressed more than predicted and the other strain repressed less. This indicates there are likely other cellular parameters that influence gene expression levels that are not included in the model. Currently the model cannot take into account variation in the protein structure of the transcription factors or the RNAP and its sigma factors. Changes in these proteins could account for some of the discrepancies between the model and the observed levels of regulation. It is likely that some global parameters that modulate transcriptional outputs which are not accounted for in the model also contribute to the disagreement with model predictions. We note that repression is a measurement of expression relative to expression in the absence of the repressor. This definition enables us to isolate the role of a particular transcription factor in regulation. Therefore, as discussed in section S8, some global regulatory parameters such as ribosomal binding sites of the relevant genes and variables such as the ribosome copy number should not impact repression levels.

From an evolutionary perspective, it is interesting that the regulation seems to be more sensitive to changes in the activator binding energy than to the activator protein copy number, as shown in Figure 3. This result might be attributed to the nature of this transcription factor. CRP is known to be a “global” transcription factor that regulates *>*50% of the *E. coli* transcription units [50]. Given its important global role in the structure of the transcriptome, changing the copy number of CRP would have a global impact on expression whereas tuning its binding affinity at a particular regulatory region has a local impact on one promoter. The regulatory knob of CRP copy number not influencing expression at the *lac* operon indicates this regulatory region may have evolved to be robust against changes in this global regulatory parameter.

The fact that the *O*_3_ operator has the possibility to change in both directions (greater or lower affinity) as reflected in Figure 4(b) suggests plasticity of the operon, allowing it to evolve according to environmental conditions. In fact this parameter changed the most among the related microbial species as shown in Figure 8(b), having species such as *Citrobacter koseri* with an operator predicted to be 5 *k_B_T* stronger than the reference value, and other species such as *Salmonella bongori* that completely lost this binding site. Although we do not yet know whether these regulatory predictions will be borne out in experimental measurements, this analysis demonstrates the utility of our sequence-to-phenotype map in interpreting the consequences of variability within the regulatory regions of sequenced genomes.

To the best of our knowledge Figure 5 shows the first quantification of how easily regulation can change given one or two point mutations along the entire promoter region. Previous studies were limited to a subset of base pairs in the Lac repressor operators and two amino acid substitutions in the Lac repressor [51]. The distribution of predicted phenotypes is very sharp close to the reference value, as a consequence the majority of the possible mutations would not be selected on. But given that regulation can change by an order of magnitude or more in both directions (increased or decreased repression) with only two mutations, changing the regulatory region of the gene could function as a fast response strategy of adaptation.

It is known from previous work that *lac* operon expression can have an impact on cell fitness [20, 21, 22, 24]. Under laboratory conditions, high expression of the *lac* operon resulted in loss of fitness due to expression of *lacY*, a transporter which imports lactose into the cell. This would suggest regulation is essential to avoid the negative consequences of *lacY* overexpression, and tight regulation would be selected. However it is possible that natural selection would act also to modulate the magnitude of the response. Strains exposed to environments with periodical bursts of lactose could trigger instantly a high gene dosage, resulting in a steeper slope on an induction curve, while strains rarely exposed to lactose would have a moderate response, i.e. a less steep induction curve. Our exploration and prediction of regulatory phenotypes in sequenced genomes shows that the biggest changes in regulation were found to increase repression (Figure 6(c)), suggesting that lactose might not be present regularly in the natural environment of some strains.

The combination of thermodynamic models with *sort-seq* generated energy matrices presented here promises to be an useful tool to study the evolution of gene regulation. This theoretical framework allows us to explore the effect that the modification of control parameters can have on the expression levels, and to predict how point mutations in gene promoter regions enable cells to evolve their gene regulatory circuits.

## 4. Materials and methods

### 4.1. Growth conditions

Unless otherwise indicated, all experiments started by inoculating the strains from frozen stocks kept at -80*^◦^*C. Cultures were grown overnight in Luria Broth (EMD, Gibbstown, NJ) at 37*^◦^*C with shaking at 250 rpm. In all of the experiments these cultures were used to inoculate three replicates for each of the relevant conditions, diluting them 1:3000 into 3 mL of M9 buffer (2 mM *MgSO*_4_, 0.10 mM *CaCl*_2_, 48 mM *Na*_2_*HPO*_4_, 22 mM *KH*_2_*PO*_4_, 8.6 mM *NaCl*, 19 mM *NH*_4_*Cl*) with 0.5% glucose and 0.2% casamino acids (here referred to as “supplemented M9”). Cells were cultured at 37*^◦^*C with shaking at 250 rpm and harvested at the indicated *OD*_600_.

### 4.2. Gene expression measurements

To perform the LacZ assay we followed the protocol used by Garcia and Phillips [26]. Strains were grown in supplemented M9 for approximately 10 generations and harvested at an *OD*_600_ around 0.4. A volume of the cells was added to Z-buffer (60 mM *Na*_2_*HPO*_4_, 40 mM *NaH*_2_*PO*_4_, 10 mM *KCl*, 1 mM *MgSO*_4_, 50 mM *β*-mercaptoethanol, pH 7.0) for a total volume of 1 mL. For fully induced cells we used 50 *μL* and for uninduced cultures we concentrated the cells by spinning down 1 mL of culture and resuspending in Z-buffer. The cells were lysed by adding 25 *μL* of 0.1% SDS and 50 *μL* of chloroform and vortexing for 15 seconds. To obtain the readout, we added 200 *μL* of 4 mg/mL 2-nitrophenyl *β*-D-galactopiranoside (ONPG). Once the solution became noticeably yellow, we stopped the reaction by adding 200 *μL* of 2.5 M *Na*_2_*CO*_3_.

To remove cell debris we spun down the tubes at 13000 × *g* for 3 minutes. 200 *μL* of the supernatant were read at *OD*_420_ and *OD*_550_ on a microplate reader (Tecan Safire2).

The absolute activity of LacZ was measured in Miller units as

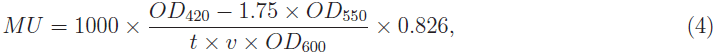

where *t* is the time we let the reaction run and *v* is the volume of cells used in mL.

The factor of 0.826 adjusts for the concentration of ONP relative to the standard LacZ assay.

### 4.3. *Measuring* in-vivo lac *repressor copy number*

To measure the repressor copy number of the natural isolates we followed the same procedure reported by Garcia and Phillips [26]. Strains were grown in 3 mL of supplemented M9 until they reached an *OD*_600_ *≈* 0.4 *–* 0.6. Then they were transferred into 47 mL of warm media and grown at 37*^◦^*C to an *OD*_600_ of 0.4–0.6. 45 mL of culture were spun down at 6000*×*g and resuspended into 900 *μ*L of breaking buffer (0.2 M Tris-*HCl*, 0.2 M *KCl*, 0.01 M Magnesium acetate, 5% glucose, 0.3 mM DTT, 50 mg/100 mL lysozyme, 50*μ*g/L phenylmethanesulfonylfluoride (PMSF), pH 7.6).

Cells were lysed by performing four freeze-thaw cycles, adding 4 *μ*L of a 2,000 Kunitz/mL DNase solution and 40 *μ*L of a 1 M *MgCl*_2_ solution and incubating at 4*^◦^*C with mixing for 4 hours after the first cycle. After the final cycle, cells were spun down at 13,000*×*g for 45 min at 4*^◦^*C. We then obtained the supernatant and measured its volume. The pellet was resuspended in 900 *μ*L of breaking buffer and again spun down at 15,000*×*g for 45 min at 4*^◦^*C. In order to review the quality of the lysing process, 2 *μ*L of this resuspended pellet was used as a control to ensure the luminescent signal of the resuspension was *<*30% of the sample.

To perform the immuno-blot we pre-wet a nitrocellulose membrane (0.2 *μ*M, Bio-Rad) in TBS buffer (20 mM *Tris – HCl*, 500 mM *NaCl*) and left it to air dry. For the standard curve a purified stock of Lac repressor tetramer [46] was serially diluted into HG105 (Δ*lacI* strain) lysate. 2 *μ*L were spotted for each of the references and each of the samples. After the samples were visibly dried the membrane was blocked using TBST (20 mM Tris Base, 140 mM *NaCl*, 0.1% Tween 20, pH 7.6) +2% BSA +5% dry milk for 1 h at room temperature with mixing. We then incubated the membrane in a 1:1000 dilution of anti-*LacI* monoclonal antibody (from mouse; Millipore) in blocking solution for 1.5 h at room temperature with mixing. The membrane was gently washed with TBS *≈* 5 times. To obtain the luminescent signal the membrane was incubated in a 1:2000 dilution of HRP-linked anti-mouse secondary antibody (GE Healthcare) for 1.5 h at room temperature with mixing and washed again *≈* 5 times with TBS. The membrane was dried and developed with Thermo Scientific Super-Signal West Femto Substrate and imaged in a Bio-Rad VersaDoc 3000 system.

### 4.4. *Constructing the* in-vivo lac *repressor energy matrix*

The energy matrix was inferred from *sort-seq* data in a manner analogous to methods described in Kinney PNAS 2010 [12]. Briefly, a library of mutant *lac* promoters was constructed in which the region [-100:25] (where coordinates are with respect to the transcription start site) was mutagenized with a 3% mutation rate. The transcriptional activity of each mutant promoter was measured by flow cytometry using a GFP reporter. To fit the LacI energy matrix, we used a Markov chain Monte Carlo algorithm to fit an energy matrix to the LacI *O*_1_ binding site by maximizing the mutual information between energies predicted by the matrix and flow cytometry measurements. The justification for maximizing mutual information is described in detail in [12, 52].

## Acknowledgments

We would like to acknowledge Ron Milo, Niv Antonovsky, Adrian Jinich, Sushant Sundaresh, Joanna Robaszewski and Hernan Garcia for useful discussions. We are grateful to Valeria Souza (UNAM) for her kind donation of the *E. coli* strains. This work was supported by the National Institutes of Health, grant numbers DP1 OD000217A (Directors Pioneer Award), R01 GM085286 and R01 GM085286B (www.nih.gov). This work was also supported by the Donna and Benjamin M. Rosen Center for Bioengineering at Caltech. The funders had no role in study design, data collection and analysis, decision to publish, or preparation of the manuscript.

## Supplemental Material

### S1. Alignment of promoter sequences

Figure S1 shows the alignment of the promoter regions of the *E. coli* wild isolates sequenced.

**Figure S1.**
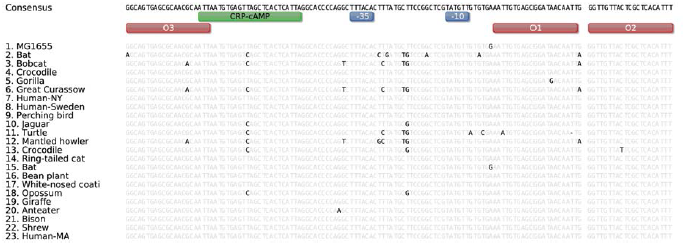
Promoter alignment of the sequenced strains. Highlighted bases differ from the consensus sequence on top. Colored boxes indicate the relevant binding sites for the Lac repressor (red), CRP (green) and RNAP (blue)

### S2. 16S rRNA sequences

To confirm the identity of the strains we analyzed 490 bp of the 16S rRNA. Figure S2 shows a schematic representation of the sequences. Colored basepairs represent mutations with respect to the consensus sequence. All sequences were found to be *≥*99% similar to the reference *E. coli MG1655* sequence.

**Figure S2.**
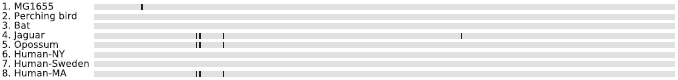
16S sequence alignment. Black lines represent mutations with respect to the consensus sequence.

### S3. Model parameters

Table S1 shows the values of the reference parameters for MG1655 obtained from different sources.

**Table S1.** Reference parameters for the strain MG1655.

| Parameter | Symbol | Value | Units | Reference |
| --- | --- | --- | --- | --- |
| $O_1$ repressor operator binding energy | $\Delta\epsilon_r^{O_1}$ | -15.3 | $k_B T$ | [1] |
| $O_2$ repressor operator binding energy | $\Delta\epsilon_r^{O_2}$ | -13.9 | $k_B T$ | [1] |
| $O_3$ repressor operator binding energy | $\Delta\epsilon_r^{O_3}$ | -9.7 | $k_B T$ | [1] |
| Repressor copy number | $R$ | 20 | tetramer/cell | Measured |
| Activator binding energy | $\Delta\epsilon_a$ | -13 | $k_B T$ | [2, 3] |
| Number of active activators | $A$ | 55 | active molecules/cell | [2] |
| RNAP binding energy for the *lac* promoter | $\Delta\epsilon_p$ | -5.35 | $k_B T$ | [4] |
| RNAP copy number | $P$ | 5500 | active molecules/cell | [5] |
| Number of nonspecific binding sites | $N_{NS}$ | $4.6 \times 10^6$ | - | GenBank: U00096.2 |
| Looping free energy between $O_1 - O_2$ | $\Delta F_{loop(l_{12})}$ | 4.7 | $k_B T$ | Fit to data from [6, 7] |
| Looping free energy between $O_1 - O_3$ | $\Delta F_{loop(l_{13})}$ | 9 | $k_B T$ | [8] |
| Looping free energy between $O_2 - O_3$ | $\Delta F_{loop(l_{23})}$ | 5.2 | $k_B T$ | Fit to data from [6, 7] |
| RNAP-CRP interaction energy | $\Delta\epsilon_{ap}$ | -5.3 | $k_B T$ | [9, 2] |
| Lac repressor - CRP interaction energy | $\Delta\epsilon_{ar}$ | -5.5 | $k_B T$ | Fit to data from [6, 7] |

### S4. Derivation of the repression level equation

Thermodynamic models of gene regulation consider that the gene expression level is proportional to the probability of finding the RNAP bound to the promoter region [3, 10, 11, 12]. This biologically simplistic but powerful predictive tool allows us to study the effect of different transcription factors in different promoter architectures. In the case of the wild-type (WT) *lac* operon promoter architecture, where we have two different transcription factors involved in the regulation - the activator CRP and the Lac repressor.

The Lac repressor molecule, when bound to the main operator *O*_1_, blocks the polymerase from binding to the promoter region, stopping the transcription of the operon. CRP plays a double role in the regulation of the operon, activating transcription by recruiting RNAP to the promoter region, and as several experiments have shown, enhancing repression by facilitating the formation of the upstream loop between the *O*_1_ – *O*_3_ operators [13, 14, 15]. Enhanced repression by CRP is due to pre-bending the DNA between 90*^◦^* and 120*^◦^* [16], thereby increasing the probability of looping by bringing the *lac* operators closer together. The model captures this effect by adding an interaction term Δ*ε_ar_* in the states where CRP is bound and the Lac repressor forms a loop between operators *O*_1_ and *O*_3_.

Assuming quasi-equilibrium conditions for the relevant processes involved in transcription, we can use the Boltzmann distribution to compute the probability of finding the RNAP bound to the promoter region, obtaining

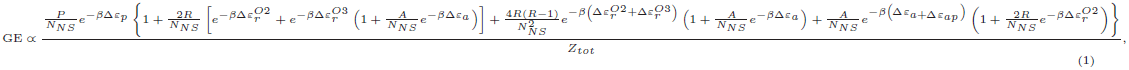

where GE stands for gene expression, *Z*_*tot*_ represents the partition function for the states shown in Figure 2 in the main text. The presence of CRP in the promoter region is not assumed to influence the kinetics of promoter escape, only the probability of RNAP binding. Tagami and Aiba [17] found that the role of CRP in the activation of the *lac* operon is restricted to the steps up to the formation of the open complex, in other words, the interaction between CRP and the RNAP are not essential for transcription after the formation of the open complex. In our model we capture this effect by including an interaction energy between CRP and the RNAP, Δ*ε*_*ap*_, that has been measured experimentally [2, 9].

In the activation mechanism proposed by Tagami and Aiba [17] CRP bends the DNA and RNAP recognizes the CRP-DNA bent complex. This model would imply that RNAP makes additional contacts with the upstream region of the promoter. Based on this model we assume that the presence of the Lac repressor bound on the *O*_3_ operator and CRP bound on its binding site (without forming a DNA loop between *O*_1_ – *O*_3_) allows transcription to occur. Since the RNAP cannot contact the upstream region of the promoter because of the presence of the repressor, the interaction energy between CRP and RNAP is not taken into account in these states.

In order to quantify the influence of Lac repressor on expression levels, we measure repression, which is the fold change in gene expression as a result of the presence of the repressor. This metric has the benefit of normalizing to a strain with an identical genetic background, thus isolating the role of the repressor in regulation. This relative measurement is defined as 

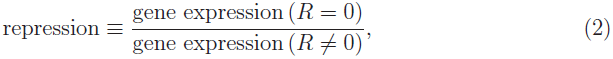

where *R* is the Lac repressor copy number. Computing this we obtain

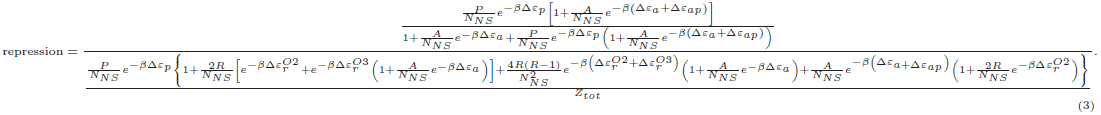

This can be further simplified, resulting in

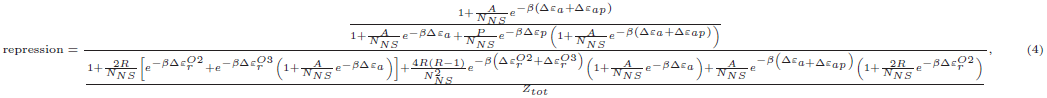

the expression we use to predict the repression level of the natural isolates.

#### S4.1. Estimating the number of active CRP molecules

The Catabolite Activator Protein, also known as cAMP-receptor protein (CRP) is a global transcriptional regulator in *E. coli* [18]. As it exists in two forms, the cAMP-CRP complex which is considered as the active state and the inactive state without cAMP bound, the number of active molecules is a function of the cAMP cellular concentration. From a thermodynamic perspective we can estimate this number as 

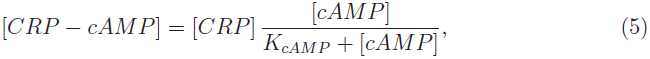

where [*CRP - cAMP*] is the concentration of active proteins, [*CRP*] is the total concentration of this transcription factor, [*cAMP*] is the cellular concentration of cAMP and *K*_*cAMP*_ is the in vivo dissociation constant of the cAMP-CRP complex.

Kuhlman et al. [2] reported the values for the CRP concentration ([*CRP*] *≈* 1500 nM) and the dissociation constant (*K*_*cAMP*_ = 10 **μ**M). Epstein et al. [19] measured the intracellular cAMP concentration in different media, including minimal media with glucose and casamino acids ([*cAMP*] *≈* 0.38*μ*M). Using these values we calculate the number of active CRP molecules as

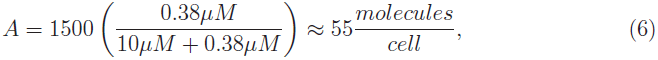

where we used the rule of thumb that 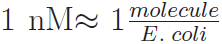. This rule of thumb is enough for our predictions since the repression level is predicted to be largely insensitive to the activator copy number as shown in Figure 3 in the main text.

#### S4.2. Estimating the number of available RNAP

In order to estimate the available number of RNAP molecules, we appeal to the work of Klumpp and Hwa [5] where they calculated the total number of RNAP molecules as well as the fraction of these molecules available for transcription as a function of the growth rate. Figure S3 shows the number of available RNAP as a function of the doubling cycles per hour.

**Figure S3.**
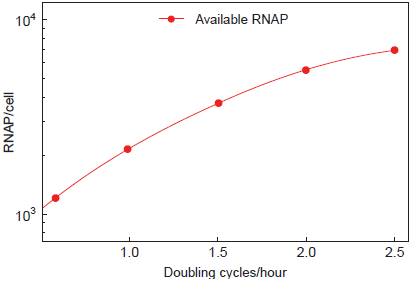
Adapted from Klumpp and Hwa [5]. RNAP available for transcription as a function of the number of doubling cycles per hour.

Using these results, we estimate 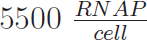 for cells grown in 0.6% glucose + 0.2% casamino acids (with a doubling time of *≈* 30 min.). We interpolate between these data to obtain the RNAP copy number for each of the natural isolates.

#### S4.3. Estimating CRP’s Binding energy

The activator binding energy was estimated as reported by Bintu et al. [3]. Using the reported dissociation constants from the specific binding site, 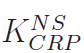, and nonspecific sequences, 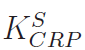, we can compute the binding energy as

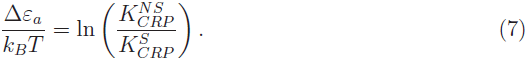

Bintu et al. also reported the following values for both dissociation constants 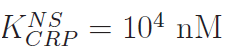 and 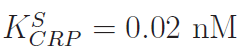 which gives us Δ*ε_a_ ≈ -*13 *k*_*B*_*T*.

### S5. Fitting parameters and testing the model

The three unknown parameters, the looping energies for the *O*_1_ – *O*_2_ and *O*_3_ - *O*_2_ loops and the decrease in the looping free energy when CRP and Lac repressor are bound at the same time, were inferred from the classic work of Oehler *et al.* [7, 6]. In these papers Oehler and collaborators measured the repression level of different *lac* operon constructs with either mutagenized or swapped Lac repressor binding sites while changing the repressor copy number. Because they reported the mutagenized sequences for the repressor binding sites we used the *sort-seq* derived energy matrix to calculate the residual energies of these modified binding sites. The three unknown parameters were fitted by minimizing the mean square error of the measurements,

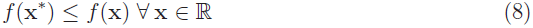

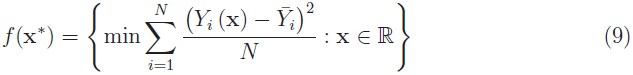

where *Y*_*i*_ is the predicted value, 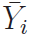 is the experimental repression level for each of the constructs measured by Oehler *et al.* and **x** are the fitting parameters. Using this method we fit for the values of Δ*F*_*loop*(*l*_13_)_, Δ*F*_*loop*(*l*_23_)_, and Δ*ε_ar_* using the data from references [7, 6]. The three parameter values are listed in Table S1.

### S6. Testing the model with different data

We used the model to predict the repression level of constructs reported by Oehler *et al.* [7, 6] and Müller *et al.* [20]. Figure S4 shows the comparison of the model predictions and the experimental results. The calculations were done using the model whose states are depicted in Figure 2, assuming a wild type repressor copy number of 10 repressors per cell, and calculating all the residual binding energies with the Lac repressor *sort-seq* derived energy matrix.

**Figure S4.**
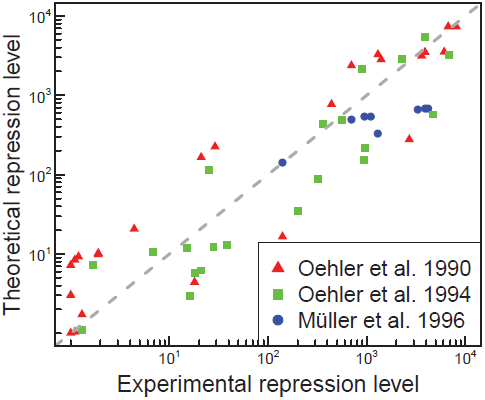
Comparing the experimental data from Oehler *et al.* [7, 6] and Müller *et al.* [20] with the model prediction.

### S7. Error propagation

To calculate a confidence interval of the model, we used the *law of error propagation* [21] where we compute the contribution of the uncertainty in parameters to the uncertainty of the repression level as

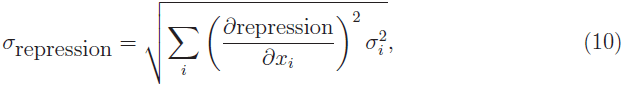

where *x*_*i*_ represents each of the parameters of the model (binding energies, transcription factors copy number, looping energies, etc.) and *σ_i_* represents the standard deviation of each of these parameters.

Paradoxically, calculating the contribution of each parameter to the uncertainty of the model requires “certainty” about the variability of these parameters. This means that we can only include the uncertainty of the parameters whose uncertainty measurements represents the natural variability in its value and not mostly error due to experimental methods. Table S2 lists the uncertainty of the parameters considered in this analysis given that the *in vivo* error was reported in the listed bibliography.

**Table S2.**
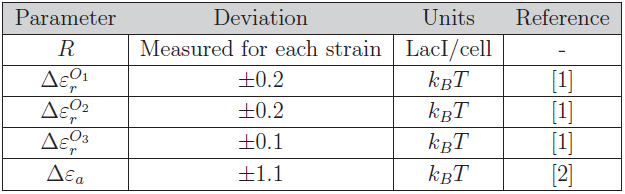
Standard deviation of the parameters considered for the calculation of the confidence interval.

We used a customized *Mathematica* script (Wolfram Research, Champaign, IL) to calculate the partial derivatives. Figure S5 reproduces Figure 7 from the main text, including the predicted standard deviation.

**Figure S5.**
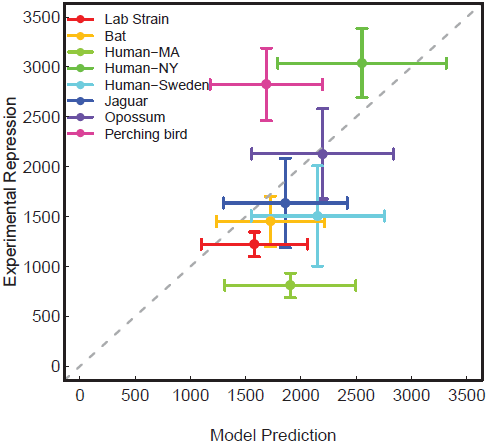
Comparison of the model prediction with the experimental measurement. Vertical error bars represent the standard deviation of at least three independent measurements each with three replicates. Horizontal error bars represent the 68% confidence interval of the model calculated by using the *law of error propagation* with the parameter uncertainties listed in Table S2.

### S8. Measuring repression level decouples growth rate effects in translation from effects in transcription

From previous work it was determined that one key regulatory parameter that is influenced by growth rate is the RNAP copy number [22]. However other cellular parameters such as ribosomal copy number and the dilution of mRNA concentration due to growth are also impacted. These parameters will influence protein copy number by influencing the efficiency of mRNA translation. In a very simple dynamical model of transcription, we can imagine that the change in the number of messenger RNA (mRNA) is proportional to the transcription rate and the degradation rate of the mRNA,

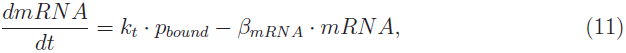

where *k*_*t*_ is the maximum transcription rate when the operon is fully induced and *p*_*bound*_ is the probability of finding the RNAP bound to the relevant promoter, as derived using statistical mechanics, *β_mRNA_* is the mRNA degradation rate and *mRNA* is the number of transcripts of the gene per cell. This equation assumes that the most relevant effect for mRNA depletion is the degradation of the transcripts, compared with the dilution effect due to the growth rate. It is known that this degradation term is not strongly affected by the growth rate [22], so we assume that this term remains constant. In steady state, when cells are in the exponential growth phase, the concentration of mRNA is

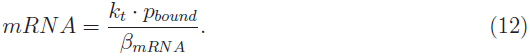

The Miller assay (LacZ assay) quantifies the level of LacZ expression, and we assume that the number of proteins is directly proportional to the mRNA copy number. Due to the relatively fast doubling time we assume that dilution is the relevant effect diminishing protein copy number, leading us to

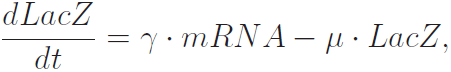

where *γ* is the proportionality constant of how many proteins per mRNA are produced, **μ** is the growth rate, and *LacZ* is the *β*-galactosidase enzyme copy number. *γ* can be a function of the growth rate due to the changes in the number of available ribosomes, but still we argue that measuring the repression level should reduce the importance of these effects. If we substitute Equation 12 into 13 and assume steady state we obtain

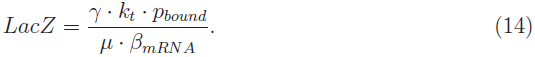

By computing the repression level as measured in the LacZ assay we obtain

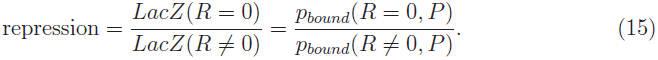

In this ratio *γ*, *k*_*t*_, **μ**, and *β_mRNA_* cancel each other leaving only a ratio of *p*_*bound*_’s.

### S9. Related microbial species *lac* operon phylogenetic tree

**Figure S6.**
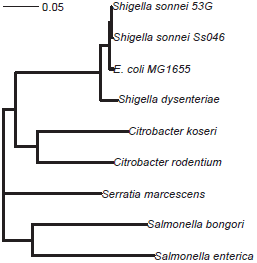
*lac* operon phylogenetic tree of diverse species with a similar *lac* promoter architecture done with the Neighbor-Joining algorithm. The scale bar represents the relative number of substitutions per sequence.

### S10. Epistasis Analysis

Epistasis can be defined as the effect of mutations on the phenotypes caused by other mutations. Our theoretical model explicitly ignores possible interactions between mutations when calculating the transcription factor binding energies with the *sort-seq* energy matrices; but the same cannot be directly assumed for the phenotypic output. As shown in Figure 3 in the main text, the phenotypic response depends on the model parameters in a highly non-linear way. Given this non-linear relation we decided to perform an epistasis analysis on the data, where we defined epistasis as [23, 24]

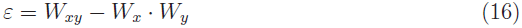

where *ε* is the epistasis, *W*_*xy*_ is the repression value for the double mutant at positions *x* and *y* normalized to the reference MG1655 repression level, and *W*_*x*_ and *W*_*y*_ are the repression values for the single mutants in their respective positions also normalized to the same reference value. This multiplicative epistasis model indicates the type of interaction between mutations; *ε* = 0 indicates no epistasis, *ε <* 0 indicates antagonistic epistasis and *ε >* 0 indicates synergistic epistasis [23].

We calculated this epistasis metric for all the double mutants of the 134 base-pairs considered in the regulatory region of the *lac* operon including the *O*_2_ downstream repressor binding site. For each pair of bases we calculated the epistasis for the two nucleotides with the biggest change with respect to our reference strain MG1655. Figure S7 shows the distribution of the epistasis values for the 8911 possible double mutants. As we initially assumed, most of the base-pairs do not interact with each other. Only 0.5% of the double mutants have an *ε <* −0.5, and 1% have an *ε >* 0.5.

**Figure S7.**
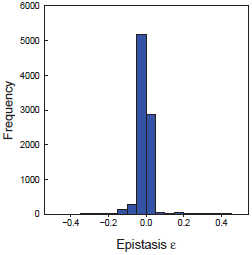
Epistasis level (Equation 16) distribution of all the possible double mutants of the *lac* operon regulatory region.

In order to find the base-pairs in the regulatory region predicted to have the biggest interactions Figure S8 shows the heat-map of the *ε* values. It is interesting to note that the few regions predicted to have significant epistasis fall mostly within a single binding site, i.e., basically no interaction is predicted between mutations located in different binding sites. The RNAP binding site is predicted to have antagonistic epistasis (*ε <* 0), while the CRP binding site is predicted to have strong synergistic epistasis (*ε >* 0). The *O*_3_ binding site also presents synergistic interactions. This predicted epistasis can be attributed to the highly non-linear dependence of the repression level on these binding energies. Since, for example, the linear regime of the *O*_1_ binding energy extends over a larger range of values (Figure 3 on the main text) two mutations are unable to move this parameter to the non-linear region and no epistasis would be expected at this binding site. Interestingly the only interactions between different binding sites are predicted to be between CRP and RNAP.

**Figure S8.**
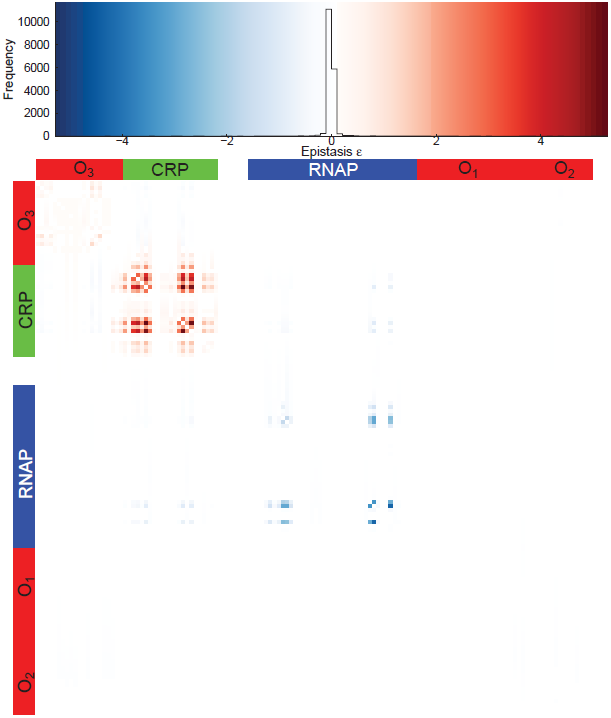
Epistasis level heat-map for all the possible double mutants. The binding sites positions are indicated with the lateral color bars.

## References

[1] Janelle R Thompson, Sarah Pacocha, Chanathip Pharino, Vanja Klepac-Ceraj, Dana E Hunt, Jennifer Benoit, Ramahi Sarma-Rupavtarm, Daniel L Distel, and Martin F Polz. Genotypic diversity within a natural coastal bacterioplankton population. Science (New York, N.Y.), 307(5713):1311–3, February 2005.

[2] Lior Zelcbuch, Niv Antonovsky, Arren Bar-Even, Ayelet Levin-Karp, Uri Barenholz, Michal Dayagi, Wolfram Liebermeister, Avi Flamholz, Elad Noor, Shira Amram, Alexander Brandis, Tasneem Bareia, Ido Yofe, Halim Jubran, and Ron Milo. Spanning high-dimensional expression space using ribosome-binding site combinatorics. Nucleic acids research, 41(9):e98, May 2013.

[3] Eran Segal and Jonathan Widom. From DNA sequence to transcriptional behaviour: a quantitative approach. Nature reviews. Genetics, 10(7):443–56, July 2009.

[4] Thomas E Hansen. The Evolution of Genetic Architecture. Annual Reviews of Ecology Evolution, and Systtematics, 37(May):123–157, 2006.

[5] Harley H McAdams, Balaji Srinivasan, and Adam P Arkin. The evolution of genetic regulatory systems in bacteria. Nature reviews. Genetics, 5(3):169–78, March 2004.

[6] J Christian Perez and Eduardo a Groisman. Evolution of transcriptional regulatory circuits in bacteria. Cell, 138(2):233–44, July 2009.

[7] G K Ackers, A D Johnson, and A M Shea. Quantitative model for gene regulation by lambda phage repressor. Proceedings of the National Academy of Sciences of the United States of America, 79(4):1129–33, February 1982.

[8] Nicolas E Buchler, Ulrich Gerland, and Terence Hwa. On schemes of combinatorial transcription logic. Proceedings of the National Academy of Sciences of the United States of America, 100(9):5136–41, April 2003.

[9] Lacramioara Bintu, Nicolas E Buchler, Hernan G Garcia, Ulrich Gerland, Terence Hwa, Jané Kondev, and Rob Phillips. Transcriptional regulation by the numbers: models. Current opinion in genetics & development, 15(2):116–24, April 2005.

[10] Lacramioara Bintu, Nicolas E Buchler, Hernan G Garcia, Ulrich Gerland, Terence Hwa, Jané Kondev, Thomas Kuhlman, and Rob Phillips. Transcriptional regulation by the numbers: applications. Current opinion in genetics & development, 15(2):125–35, April 2005.

[11] Marc S Sherman and Barak A Cohen. Thermodynamic state ensemble models of cis-regulation. PLoS computational biology, 8(3):e1002407, January 2012.

[12] Justin B Kinney, Anand Murugan, Curtis G Callan, and Edward C Cox. Using deep sequencing to characterize the biophysical mechanism of a transcriptional regulatory sequence. PNAS, 107(20):9158–63, 2010.

[13] Eilon Sharon, Yael Kalma, Ayala Sharp, Tali Raveh-Sadka, Michal Levo, Danny Zeevi, Leeat Keren, Zohar Yakhini, Adina Weinberger, and Eran Segal. Inferring gene regulatory logic from high-throughput measurements of thousands of systematically designed promoters. Nature biotechnology, 30(6):521–30, June 2012.

[14] Alexandre Melnikov, Anand Murugan, Xiaolan Zhang, Tiberiu Tesileanu, Li Wang, Peter Rogov, Soheil Feizi, Andreas Gnirke, Curtis G Callan, Justin B Kinney, Manolis Kellis, Eric S Lander, and Tarjei S Mikkelsen. Systematic dissection and optimization of inducible enhancers in human cells using a massively parallel reporter assay. Nature biotechnology, 30(3):271–7, March 2012.

[15] Robert C. Brewster, Daniel L. Jones, and Rob Phillips. Tuning Promoter Strength through RNA Polymerase Binding Site Design in *Escherichia coli*. PLoS Computational Biology, 8(12):e1002811, December 2012.

[16] C J Wilson, H Zhan, L Swint-Kruse, and K S Matthews. The lactose repressor system: paradigms for regulation, allosteric behavior and protein folding. Cellular and molecular life sciences : CMLS, 64(1):3–16, January 2007.

[17] W S Reznikoff. The lactose operon-controlling elements: a complex paradigm. Molecular microbiology, 6(17):2419–22, September 1992.

[18] Y Setty, A E Mayo, M G Surette, and U Alon. Detailed map of a cis-regulatory input function. Proceedings of the National Academy of Sciences of the United States of America, 100(13):7702– 7, June 2003.

[19] Thomas Kuhlman, Zhongge Zhang, Milton H Saier, and Terence Hwa. Combinatorial transcriptional control of the lactose operon of *Escherichia coli*. Proceedings of the National Academy of Sciences of the United States of America, 104(14):6043–8, April 2007.

[20] Antony M Dean. Selection and Neutrality in Lactose Operons of *Escherichia coli*. Genetics, 123:441–54, 1989.

[21] Erez Dekel and Uri Alon. Optimality and evolutionary tuning of the expression level of a protein. Nature, 436(7050):588–92, July 2005.

[22] Lilia Perfeito, Stephane Ghozzi, Johannes Berg, Karin Schnetz, and Michael Lässig. Nonlinear Fitness Landscape of a Molecular Pathway. PLoS genetics, 7(7):1–10, 2011.

[23] Frank J Poelwijk, Philip D Heyning, Marjon G J de Vos, Daniel J Kiviet, and Sander J Tans. Optimality and evolution of transcriptionally regulated gene expression. BMC systems biology, 5(1):128, January 2011.

[24] Matt Eames and Tanja Kortemme. Cost-benefit tradeoffs in engineered *lac* operons. Science (New York, N.Y.), 336(6083):911–5, May 2012.

[25] Jose M G Vilar. Accurate prediction of gene expression by integration of DNA sequence statistics with detailed modeling of transcription regulation. Biophysical journal, 99(8):2408–13, October 2010.

[26] Hernan G Garcia and Rob Phillips. Quantitative dissection of the simple repression input-output function. Proceedings of the National Academy of Sciences of the United States of America, 108(29):12173–8, July 2011.

[27] James Q. Boedicker, Hernan G. Garcia, and Rob Phillips. Theoretical and Experimental Dissection of DNA Loop-Mediated Repression. Physical Review Letters, 110(1):018101, January 2013.

[28] H Tagami and H Aiba. Role of CRP in transcription activation at *Escherichia coli lac* promoter: CRP is dispensable after the formation of open complex. Nucleic acids research, 23(4):599–605, February 1995.

[29] J M Hudson and M G Fried. Co-operative interactions between the catabolite gene activator protein and the lac repressor at the lactose promoter. Journal of molecular biology, 214(2):381– 96, July 1990.

[30] Valeria Souza, Martha Rocha, Aldo Valera, and Luis E Eguiarte. Genetic Structure of Natural Populations of *Escherichia coli* in Wild Hosts on Different Continents. Applied and Environmental Microbiology, 65(8):3373–3385, 1999.

[31] Eran Segal, Tali Raveh-Sadka, Mark Schroeder, Ulrich Unnerstall, and Ulrike Gaul. Predicting expression patterns from regulatory sequence in *Drosophila* segmentation. Nature, 451(7178):535–40, January 2008.

[32] Leonor Saiz and Jose M G Vilar. *Ab initio* thermodynamic modeling of distal multisite transcription regulation. Nucleic acids research, 36(3):726–31, February 2008.

[33] Jose M G Vilar and Stanislas Leibler. DNA Looping and Physical Constraints on Transcription Regulation. Journal of Molecular Biology, 331(5):981–989, August 2003.

[34] Jose M G Vilar and Leonor Saiz. DNA looping in gene regulation: from the assembly of macromolecular complexes to the control of transcriptional noise. Current opinion in genetics & development, 15(2):136–44, April 2005.

[35] Leonor Saiz and Jose M G Vilar. Multilevel deconstruction of the *In vivo* behavior of looped DNA-protein complexes. PloS one, 2(4):e355, January 2007.

[36] Leonor Saiz and Jose M G Vilar. DNA looping: the consequences and its control. Current opinion in structural biology, 16(3):344–50, June 2006.

[37] Leonor Saiz, J Miguel Rubi, and Jose M G Vilar. Inferring the *in vivo* looping properties of DNA. Proceedings of the National Academy of Sciences of the United States of America, 102(49):17642– 5, December 2005.

[38] Mattias Rydenfelt, Robert Cox, Hernan Garcia, and Rob Phillips. Statistical mechanical model of coupled transcription from multiple promoters due to transcription factor titration. Physical Review E, 89(1):012702, January 2014.

[39] H C Nelson and R T Sauer. Lambda repressor mutations that increase the affinity and specificity of operator binding. Cell, 42(2):549–58, September 1985.

[40] Johan Elf, Gene-Wei Li, and X. Sunney Xie. Probing Transcription Factor Dynamics at the Single-Molecule Level in a Living Cell. Science (New York, N.Y.), 316(5828):1191–1194, 2007.

[41] Larry J Friedman and Jeff Gelles. Mechanism of transcription initiation at an activator-dependent promoter defined by single-molecule observation. Cell, 148(4):679–89, February 2012.

[42] Stefan Oehler, Elisabeth R Eismann, Helmut Krämer, and Benno Müller-Hill. The three operators of the *lac* operon cooperate in repression. EMBO Journal, 9(4):973–979, 1990.

[43] Hans Bremer and Patrick P Dennis. Modulation of Chemical Composition and Other Parameters of the Cell by Growth Rate. Escherichia coli and Salmonella: cellular and molecular biology, (122):1553–1569, 1996.

[44] Stefan Klumpp and Terence Hwa. Growth-rate-dependent partitioning of RNA polymerases in bacteria. Proceedings of the National Academy of Sciences of the United States of America, 105(51):20245–50, December 2008.

[45] K M Vossen, D F Stickle, and M G Fried. The mechanism of CAP-*lac* repressor binding cooperativity at the *E. coli* lactose promoter. Journal of molecular biology, 255(1):44–54, January 1996.

[46] Stephanie Johnson, Martin Lindén, and Rob Phillips. Sequence dependence of transcription factor-mediated DNA looping. Nucleic acids research, 40(16):7728–38, September 2012.

[47] Stefan Oehler, Michele Amouyal, Peter Kolkhof, Brigitte Von Wilcken-Bergmann, and Benno Müller-Hill. Quality and position of the three *lac* operators of *E. coli* define efficiency of repression. EMBO Journal, 13(14):3348–3355, 1994.

[48] Frank J Poelwijk, Daniel J Kiviet, and Sander J Tans. Evolutionary potential of a duplicated repressor-operator pair: simulating pathways using mutation data. PLoS computational biology, 2(5):e58, May 2006.

[49] Moisés Santillán. On the Use of the Hill Functions in Mathematical Models of Gene Regulatory Networks. Mathematical Modelling of Natural Phenomena, 3(2):85–97, 2008.

[50] Agustino Martínez-Antonio and Julio Collado-Vides. Identifying global regulators in transcriptional regulatory networks in bacteria. Current Opinion in Microbiology, 6(5):482– 489, October 2003.

[51] Alexandre Dawid, Daniel J Kiviet, Manjunatha Kogenaru, Marjon de Vos, and Sander J Tans. Multiple peaks and reciprocal sign epistasis in an empirically determined genotype-phenotype landscape. Chaos (Woodbury, N.Y.), 20(2):026105, June 2010.

[52] Justin B Kinney, Gasper Tkacik, and Curtis G Callan. Precise physical models of protein-DNA interaction from high-throughput data. Proceedings of the National Academy of Sciences of the United States of America, 104(2):501–6, January 2007.

## References

[1] Garcia H. G and Phillips R. 2011. Quantitative dissection of the simple repression input-output function. Proceedings of the National Academy of Sciences of the United States of America, 108(29):12173–8.

[2] Kuhlman T, Zhang Z, Saier M. H, and Hwa T. 2007. Combinatorial transcriptional control of the lactose operon of *Escherichia coli*. Proceedings of the National Academy of Sciences of the United States of America, 104(14):6043–8.

[3] Bintu L, Buchler N. E, Garcia H. G, Gerland U, Hwa T, Kondev J, and Phillips R. 2005. Transcriptional regulation by the numbers: models. Current opinion in genetics & development, 15(2):116–24.

[4] Brewster R. C, Jones D. L, and Phillips R. 2012. Tuning Promoter Strength through RNA Polymerase Binding Site Design in *Escherichia coli*. PLoS Computational Biology, 8(12):e1002811.

[5] Klumpp S and Hwa T. 2008. Growth-rate-dependent partitioning of RNA polymerases in bacteria. Proceedings of the National Academy of Sciences of the United States of America, 105(51):20245– 50.

[6] Oehler S, Eismann E. R, Krämer H, and Müller-Hill B. 1990. The three operators of the *lac* operon cooperate in repression. EMBO Journal, 9(4):973–979.

[7] Oehler S, Amouyal M, Kolkhof P, Wilcken-Bergmann B. V, and Müller-Hill B. 1994. Quality and position of the three *lac* operators of *E. coli* define efficiency of repression. EMBO Journal, 13(14):3348–3355.

[8] Boedicker J. Q, Garcia H. G, and Phillips R. 2013. Theoretical and Experimental Dissection of DNA Loop-Mediated Repression. Physical Review Letters, 110(1):018101.

[9] Kinney J. B, Murugan A, Callan C. G, and Cox E. C. 2010. Using deep sequencing to characterize the biophysical mechanism of a transcriptional regulatory sequence. PNAS, 107(20):9158–63.

[10] Ackers G. K, Johnson A. D, and Shea A. M. 1982. Quantitative model for gene regulation by lambda phage repressor. Proceedings of the National Academy of Sciences of the United States of America, 79(4):1129–33.

[11] Buchler N. E, Gerland U, and Hwa T. 2003. On schemes of combinatorial transcription logic. Proceedings of the National Academy of Sciences of the United States of America, 100(9):5136– 41.

[12] Sherman M. S and Cohen B. A. 2012. Thermodynamic state ensemble models of cis-regulation. PLoS computational biology, 8(3):e1002407.

[13] Hudson J. M and Fried M. G. 1990. Co-operative interactions between the catabolite gene activator protein and the lac repressor at the lactose promoter. Journal of molecular biology, 214(2):381–96.

[14] Hudson J. M and Fried M. G. 1996. DNA looping and lac repressor-CAP interaction. Science, 82:2–3.

[15] Vossen K. M, Stickle D. F, and Fried M. G. 1996. The mechanism of CAP-*lac* repressor binding cooperativity at the *E. coli* lactose promoter. Journal of molecular biology, 255(1):44–54.

[16] Schultz S. C, Shields G. C, Steitz T. A, and Stejtz T. A. 1991. Crystal Structure of a CAP-DNA Complex: The DNA of Is Bent by 90. *Science (New York*, N.Y.), 253(5023):1001–1007.

[17] Tagami H and Aiba H. 1995. Role of CRP in transcription activation at *Escherichia coli lac* promoter: CRP is dispensable after the formation of open complex. Nucleic acids research, 23(4):599–605.

[18] Martínez-Antonio A and Collado-Vides J. 2003. Identifying global regulators in transcriptional regulatory networks in bacteria. Current Opinion in Microbiology, 6(5):482–489.

[19] Epstein W, Rothman-Denes L. B, and Hesse J. 1975. Adenosine 3’:5’-cyclic monophosphate as mediator of catabolite repression in Escherichia coli. Proceedings of the National Academy of Sciences of the United States of America, 72(6):2300–4.

[20] Müller J, Oehler S, and Müller-Hill B. 1996. Repression of lac promoter as a function of distance, phase and quality of an auxiliary lac operator. Journal of molecular biology, 257(1):21–9.

[21] Ku H. H. 1966. Notes on the Use of Propagation of Error Formulas. Journal of Research of the National Bureau of Standards, 70C(4):75–79.

[22] Klumpp S, Zhang Z, and Hwa T. 2009. Growth rate-dependent global effects on gene expression in bacteria. Cell, 139(7):1366–75.

[23] Segrè D, Deluna A, Church G. M, and Kishony R. 2005. Modular epistasis in yeast metabolism. Nature genetics, 37(1):77–83.

[24] Mani R, St Onge R. P, Hartman J. L, Giaever G, and Roth F. P. 2008. Defining genetic interaction. Proceedings of the National Academy of Sciences of the United States of America, 105(9):3461–6.

